# Alternative polyadenylation and the sex-specific gene expression program in hemp

**DOI:** 10.64898/2026.05.13.725035

**Authors:** Ashwini Shivakumar, Arthur G. Hunt, Manohar Chakrabarti

**Author notes:** Corresponding author – Manohar Chakrabarti. **Data availability statement:** The PATSeq short read sequences generated in this research are available at NCBI under Bioproject PRJNA900205. Other data and analyses can be found in the Supplemental Materials.

## Abstract

Hemp (Cannabis sativa) produces a wide array of medicinally significant compounds, including cannabidiol (CBD). These compounds are predominantly synthesized in female hemp inflorescences. The proposed research utilizes next-generation sequencing-based transcriptome analysis using a 3□-end-directed approach to identify differentially expressed genes between male and female hemp plants at the early vegetative stage. 886 differentially expressed genes (DEGs) were identified, a majority of which were upregulated in males compared to females. We hypothesized that alternative RNA processing contributes to sex-specific gene expression. To this end, 932 genes were identified that exhibited significant changes in poly(A) site usage when comparing males and females. These genes were much more likely to be differentially expressed, supportive of this hypothesis. Males tend to have longer 3’ UTRs with canonical motifs found in the Near-Upstream Elements (NUE), compared to the shorter 3’ UTRs in females, which have A-rich motifs near the cleavage site. This suggests that polyadenylation remodels hemp mRNAs with distal poly(A) sites being preferred in males. To further investigate when this sex-specific gene expression program is established, RNA was isolated from plants at various developmental stages, such as developing seeds, four-day-old seedlings, and different developmental stages up to four weeks after sowing. Diagnostic male-specific genes were analyzed using RT/PCR. The results indicate that sex-specific gene expression is not evident in seeds but rather is set during or after germination.

**Significance:**

- Hemp males tend to have longer 3’ UTRs with canonical motifs found in the Near-Upstream Elements (NUE), compared to the shorter 3’ UTRs in females, which have A-rich motifs near the cleavage site.
- The sex-specific gene expression program is not yet established in mature seed but is set in the time between germination and 4 days of growth.

## Introduction

Hemp (*Cannabis sativa*) has garnered significant attention due to its multifaceted industrial and medicinal applications for bast fiber, oilseed/grain, and high-value secondary metabolites. Industrial hemp and marijuana are both derived from the same plant species *Cannabis sativa L*. Although cannabis is one of the most chemically well studied plants, many of its chemical constituents are not well characterized for their biological activity, hence termed as ‘neglected pharmacological treasure trove’ (Mechoulam, 2005). The primary psychoactive agent in cannabinoids is Δ^9^-tetrahydrocannabinolic acid (THCA). Besides THCA, cannabis also produces non-psychoactive compound such as cannabidiol (CBD), cannabichromene (CBC), Δ^9^-tetrahydrocannabivarin (THCV), which also possess pharmacological activities (van Bakel et al., 2011, Izzo et al., 2009, Mechoulam, 2005). Among these cannabinoids, THCA and CBDA (cannabidiolic acid) are structural isomers derived from the same precursor, cannabigerolic acid (CBGA), and are synthesized by the same pathway, in which the final step is catalyzed by THCA synthase and CBDA synthase. These acidic forms subsequently undergo decarboxylation with heat/light to yield CBD and THC, the principal bioactive compounds of commercial interest (Toth et al., 2020). (Marks et al., 2009). In the United States, the 2018 Agriculture Improvement Act (Farm Bill) distinguished hemp from marijuana by defining hemp as *C. sativa* with Δ^9^ THC <0.3% on a dry-weight basis, enabling regulated production and catalyzing the rapid expansion of hemp supply chains (FDA 2019, July 25, Hemp Production and the 2018 Farm Bill).

Cannabidiol (CBD) possesses important medicinal properties. It was reported to be anti-epileptic as it was implicated in reducing the frequency and severity of seizures (Rosenberg et al., 2015). It has also been highlighted as a potential analgesic, antidepressant, and anti-inflammatory compound (Fusar-Poli et al., 2009, Burstein et al., 1984, Burstein, 2015). CBD has been used as a therapeutic agent for breast cancer (Elbaz et al., 2015). Additionally, officially registered clinical trials of CBD for the treatment of schizophrenia were performed (Hagerty et al., 2015). Apart from its therapeutic benefits, the fiber applications and nutritional qualities of hemp make it significant for human health. Strong hemp fibers are used to manufacture construction materials, papers, and textiles. Hemp seeds are excellent grains because they are high in protein, good fats, and minerals. Different hemp products have different market values, and the demand for CBD products, natural fibers, and textiles made from hemp is driving the rapid expansion of the global hemp market (Andre et al., 2016).

*Cannabis* is an annual herb with a predominantly dioecious breeding system, and the cannabinoid composition and content vary greatly among different cultivars. A high THC/low CBD chemotype is regarded as a drug’ type or ‘marijuana, while a low THC/high CBD chemotype is termed as ‘hemp’(van Bakel et al., 2011). Cannabinoids, such as THC and CBD, are formed in the capitate-stalked glandular trichomes on the inflorescence bracts, and female plants generally show higher trichome densities and greater cannabinoid accumulation than males. Additionally, pollination and fertilization of female flowers typically lower phyto-cannabinoid levels relative to unpollinated females; therefore, maximal cannabinoid production is achieved in unfertilized inflorescences (Alberti et al., 2025, Hazekamp, 2009). Consistent with this, industry-oriented genomic studies have emphasized that CBD yields are maximized in unpollinated female hemp plants (Toth et al., 2020). Beyond cannabinoid production, the reproductive biology of hemp also influences agronomic traits, such as fiber quality and seed yield. Comparative evaluations have shown pronounced differences between dioecious and monoecious hemp accessions in bast fiber amount and composition, as well as sex-linked contrasts in lignification, such as females generally having more lignified fiber, while males produce finer fibers (Petit et al., 2020a). Thus, the sex determination of cannabis plants is of significant economic importance and has been an active research focus (Hirata, 1927, Divashuk et al., 2014). From an agronomic standpoint, it is of great value if female hemp plants can be detected early in the growing season, as this will drastically reduce the costs associated with various agricultural inputs. Thus, sex determination in hemp is of paramount importance to the hemp industry.

To operationalize female-biased production or identify sex before flowering, growers currently rely on two strategies with practical tradeoffs. First, genetic testing: sex-specific genetic regions in hemp plants were identified and used to develop DNA-based molecular markers to determine the sex of hemp plants early on, even before they begin to form reproductive structures. However, this approach requires a well-equipped molecular biology laboratory for genotyping and considerable expertise in molecular biology; moreover, its performance can vary by population and genetic background (Toth et al., 2020). Second, chemical or hormonal treatment: ‘Feminized’ hemp seeds can be produced by treating hemp plants with silver salts (silver thiosulphate or silver nitrate) or phytohormones. While this approach is effective across medicinal, grain, and fiber types, silver nitrate is phytotoxic and outcomes can be genotype and protocol dependent and this method is also time consuming, inconsistent, and expensive (Baek and Vergara, 2025) (Petit et al., 2020b). While DNA markers can determine sex prior to the reproductive stage, they do not reveal the regulatory mechanisms underlying sex determination (Petit et al., 2020a, Pandey et al., 2025). A transcriptome approach addresses this gap by directly assaying the regulatory state of young and mature tissues. RNA-seq data detect sex-biased expression prior to full floral differentiation and nominate plausible X/Y-linked transcriptional regulators (Shi et al., 2025). Comparable evidence in other dioecious plants such as *Mercurialis annua* (Cossard et al., 2019)*, Salix paraplesia (Cai et al., 2021), Spinacia oleracea* (Li et al., 2020)*, Silene latifolia* (Zluvova et al., 2010) and *Pistacia chinensis* (Xiong et al., 2013) show that expression signatures can separate the sexes early, providing a foundation for deployable biomarkers.

The objective of the proposed research was to identify genes that are differentially expressed in male and female hemp plants at the early vegetative stage and to examine how alternative RNA processing contributes to sex-specific gene expression. Differentially expressed genes in male and female can be utilized to develop easily deployable biomarkers to distinguish sex in the early developmental stages of hemp plants or to generate female-only hemp lines, which can save a number of resources, including space, water, agricultural inputs, and time. In addition, understanding alternative RNA processing may provide insights into the molecular mechanisms that regulate sex-specific development in hemp. Our project hypothesized that genetic regulation and alternative polyadenylation (APA) regulation of early vegetative development differ between the sexes of hemp plants. Based on this hypothesis, the objective of the current project was to employ a next-generation sequencing-based 3′-end-directed transcriptome approach to identify DEGs and APA-regulated genes at the early vegetative stage. The resulting differentially expressed candidates are positioned as practical early biomarkers for future manipulation of sex pathways, for example, the development of female-only lines by silencing male sex-specific or developmentally essential transcript isoforms with modern genome editing strategies such as CRISPR-Cas9.

## Results

### Identification of genes whose sex-specific expression is apparent in young vegetative tissues

The goal of this study was to identify hemp genes whose sex-specific expression is apparent in young plants, well before sexual development is manifested. To this end, we grew 40 hemp plants (Finola cultivar) from seeds and monitored them until reproductive maturity to determine their sex phenotypically. Of the 40 plants, two did not survive, leaving 38 plants. At the reproductive maturity stage, we observed 15 male and 23 female plants. RNA was isolated from these plants and used in subsequent transcriptomics experiment.

Transcriptome analysis was conducted using a 3□-end sequencing method. This approach (abbreviated as PATSeq) entails the production, sequencing, and analysis of short cDNAs that query the mRNA-poly(A) junction; these sequences can be used to quantify genome-wide gene expression and to assess the genome-wide usage of poly(A) sites. PATseq libraries were produced from six males and six females and sequenced on the Illumina platform. Raw reads were mapped to a set of organellar and ribosomal RNA references to remove reads that map to these regions, and the remaining reads demultiplexed, trimmed, and mapped the hemp genome (*Cannabis sativa* cultivar Finola). Mapped reads were used for analysis of poly(A) site usage and overall gene expression as described in the Methods.

Principal Component Analysis (PCA) was conducted to evaluate transcriptional variability among replicates and between the two sexes. The PCA plot showed a clear separation between male and female plants in terms of their gene expression profiles (Figure 1A). Hierarchical clustering using gene expression values was performed to assess variations within and between samples. This analysis revealed clustering of replicates for each sex (Figure 1B), further supporting the reproducibility of our experiment. Subsequently, we analyzed differentially expressed genes (DEGs). This analysis identified 886 DEGs (Figure 2A, Figure S1 and Data S2). Male- and female-specific genes were broadly distributed across the genome (Figure 2B).

**Figure 1.**
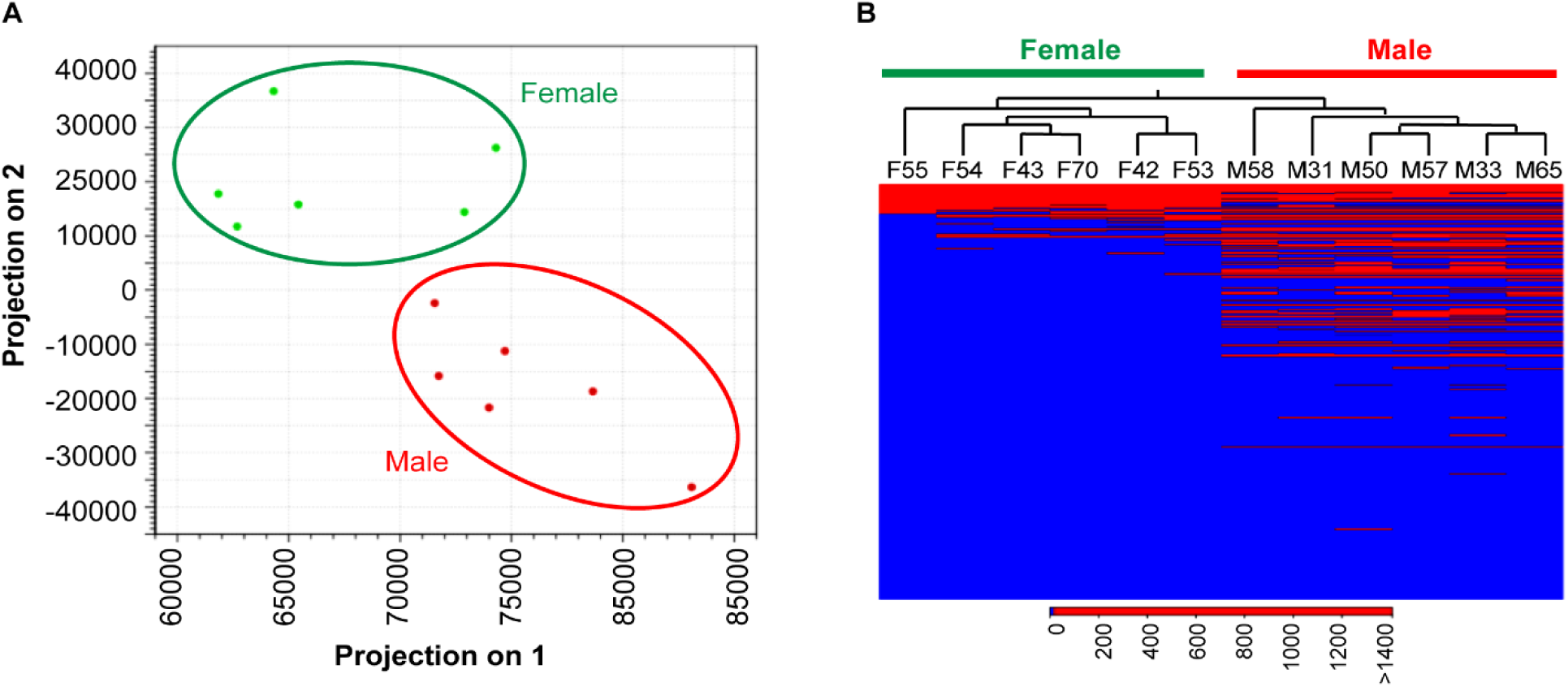
Clustering for male and female hemp samples. (A) Principal component analysis and (B) Hierarchical clustering of samples from both sexes. Only DEGs were used for the analysis and LOG_2_-transformed values were used for the analysis. Green and red color represent female and male samples, respectively. (B) Columns represent samples (present on the top) grouped by sex (green: female; red: male), and rows represent features showing differential expression patterns The scale represent log_2_-transformed values of normalized expression.

**Figure 2.**
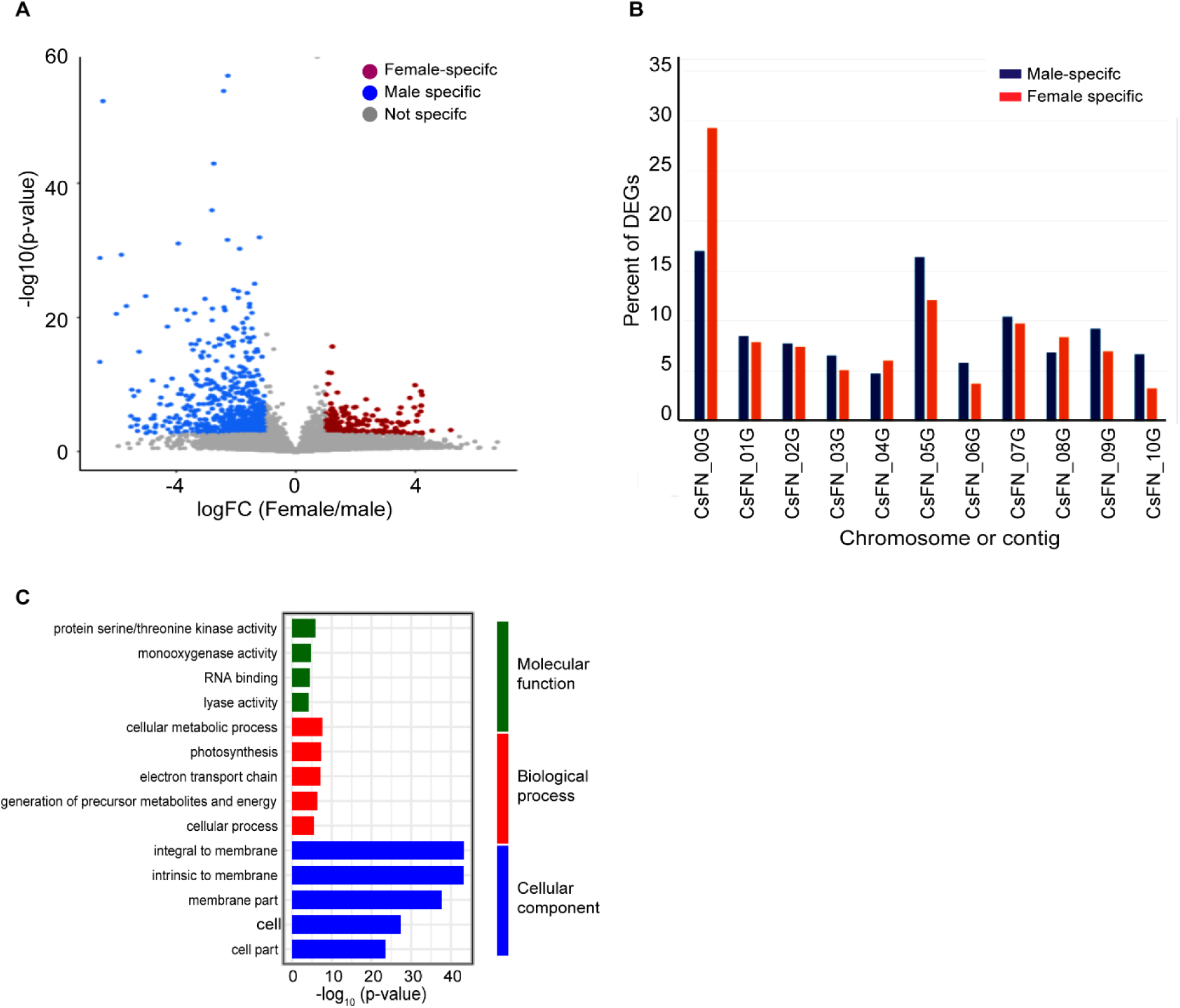
Differential gene analsyis of the male and female hemp. (A) Volcano plot showing differentially expressed genes (DEGs) between male and female samples. (B) bar plot showing the percent od DEGs spred across finnola chromosome. (C) bar plot illustrating the top 5 enriched Gene Ontology (GO) terms from each catogery (molecular function, biolofgical process and cellular compoment) for the DEGs. In panel (A), the X-axis represents log_₂_-transformed fold changes (Male/Female) calculated from normalized gene expression values, and the Y-axis shows the −log_₁₀_-transformed *p*-values. Grey dots indicate non-differentially expressed genes, red and blue dots denote significantly differentially expressed genes. In panel (C), X-axis shows the −log_₁₀_-transformed *p*-values of GO term, and the color indicating the category (molecular function, biolofgical process and cellular compoment).

A Gene Ontology (GO) analysis was performed to identify pathways enriched among the male- and female-specific DEGs (Figure 2C, Data S3). This analysis revealed a significant enrichment of genes associated with protein kinase activity, RNA binding, monooxygenase activity, and lyase activity within the molecular function category; cellular metabolic processes, photosynthesis, and electron transport chain within the biological processes category; and integral and intrinsic membrane within the cellular component category. These enrichments align with previous findings in plant reproductive biology. For instance, several RNA-binding proteins are highly expressed during floral development and stress response in plants (Lorković, 2009, Pfaff et al., 2018). Monooxygenases such as CYP703A2-A and CYP703A2-D are crucial for sporopollenin formation and pollen fertility in cotton (*Gossypium hirsutum*) (Ma et al., 2022a). Similarly, GO terms such as binding, catalytic activity, and cellular and metabolic processes are associated with pollen fertility and anther development in *Lycium barbarum* (Zhang et al., 2024).

### Sex-specific gene expression is not in place early in seed development

To further confirm the results of the differential gene expression analysis, RT-PCR was performed. For this, three genes differentially expressed between the two sexes (CsFN_10G0001520, CsFN_00G0021400, and CsFN_00G0044120) and one gene (CsFN_00G0039430) that did not display differential expression were selected from the DEG analysis. RT-PCR analysis was conducted using samples from six plants (three males and three females) that were not used for the PATSeq study. The RT-PCR results confirmed the DEG analysis findings. The three genes identified as male-specific showed expression in the three male samples but not in the female samples (Figure 3). In addition, the expression of the constitutive gene was apparent in all six samples.

**Figure 3.**
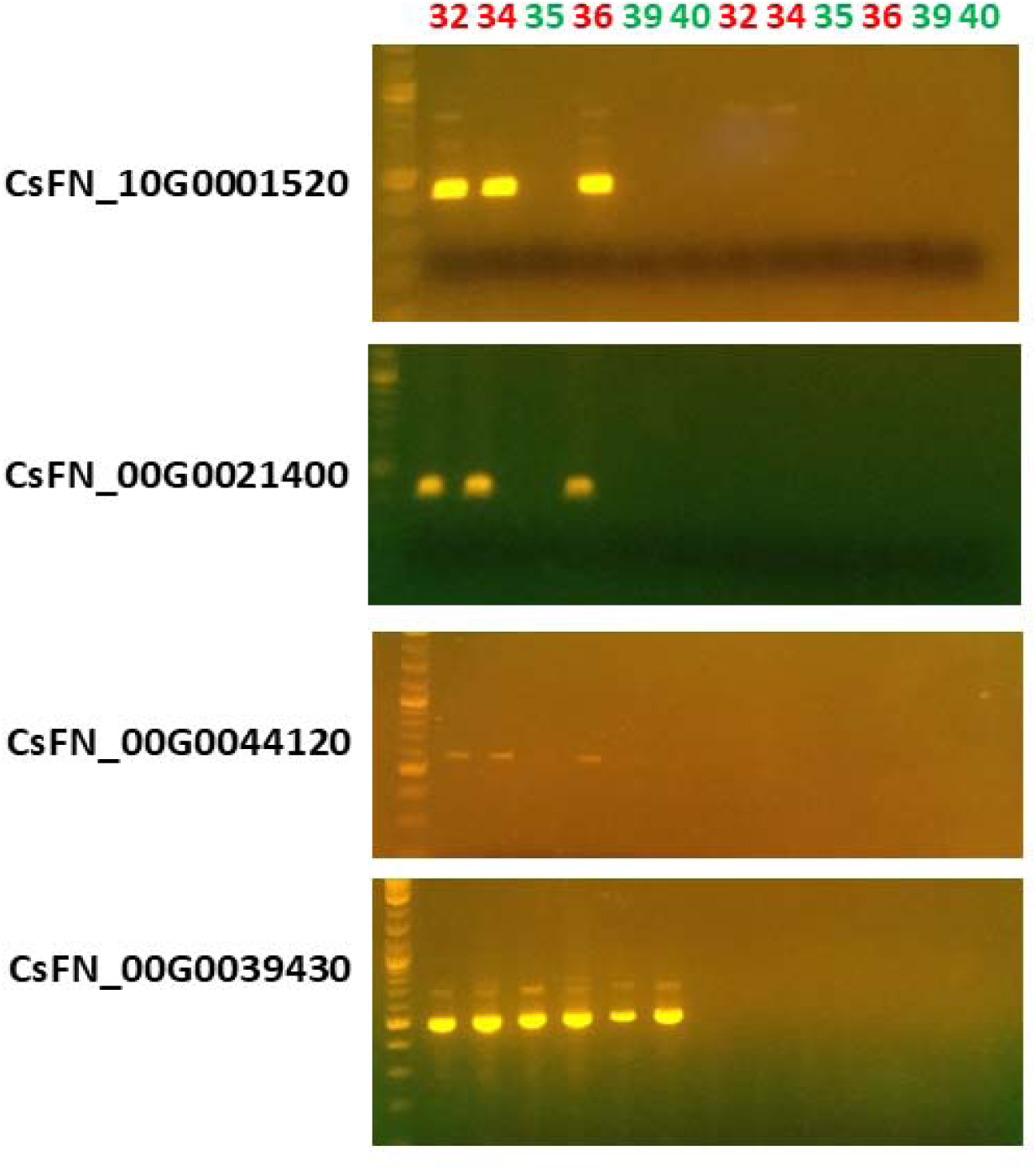
RT-PCR analysis to confirm differentially expressed genes selected by the transcriptome analysis in finola cultivar. Lane-1 represents DNA ladder, Lane-2 to 7 and 8 to 13 represent PCR products of RT and no-RT reactions of sample - 32, 34, 35, 36, 39, and 40, respectively. Red and green color represent males and females, respectively.

DEG analyses were performed using RNA isolated from the Finola cultivar. To test whether the identification of male-specific genes can be generalized to other cultivars, an analogous RT/PCR experiment was conducted using RNA isolated from 4-day old plants of the X-59 hemp cultivar. As shown in Figure 4, the expression of genes identified as male-specific in the Finola cultivar was also male-specific in the X-59 cultivar (Figure 4A). These genes were used to further explore the trajectory of sex-specific gene expression throughout the growth of hemp plants. RNA was isolated from males and females at 4, 7, 14, 21, and 28 days of growth. RT-PCR conducted on these various samples showed a consistent pattern, wherein expression was observed in males across this time frame, but not in females (Figure 4A). This result indicates that the male-specific gene expression program was established before the 4-day time point. To explore this further, RNA was isolated from individual hemp seed embryo and analyzed by RT/PCR. The results indicated that the expression of male-specific genes varied from seed to seed. Whereas all 4-day old (or older) males showed consistent expression of the three marker genes used in this study, and all such females showed no expression, individual seeds were found to express a variable number of these markers (Figure 4B). These results suggest that the sex-specific expression program is not yet established in mature seeds.

**Figure 4.**
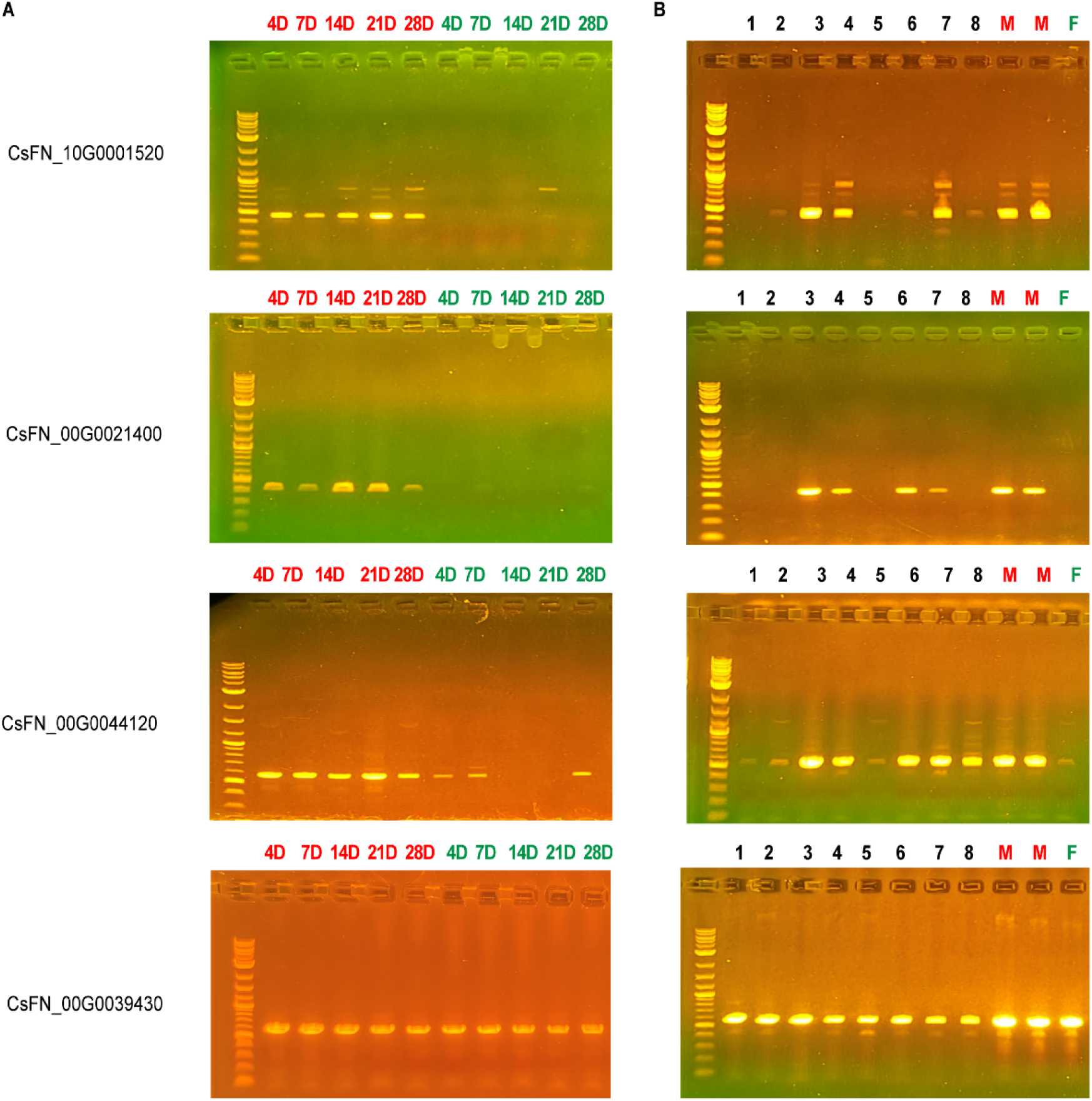
RT-PCR analysis validating the selected differential expression genes in cultivar X-59. (A) Expression patterns across developmental stages and (B) in individual developing seed embryo. Lane 1 represents the DNA ladder. In panel (A) lanes 1–5 correspond to RT-PCR products from male plants, and lanes 6–10 from female plants, collected at 4, 7, 14, 21, and 28 days of growth, respectively. Panel (B), lanes 1–8 correspond to RT-PCR products from embryos of developing seeds. “M” denotes male samples and “F” denotes female samples. D denotes for sample collected from days after sowing. Red and green color represent males and females, respectively.

### Polyadenylation site choice in male and female plants

To assess poly(A) site usage in males and females and ascertain whether differences might exist, PATSeq reads were used to identify poly(A) sites that were then grouped into clusters of adjacent sites (poly (A) site Clusters, or PACs). A total of 69,596 PACs mapped to 18,618 genes (Table 1, Data S4). As observed in other plants, the majority (>76%) of genes possessed more than one PAC (Figure 5A). A total of 1290 PACs showed significant differences in usage between males and females; these PACs defined a set of 932 genes whose poly(A) site profiles were different between males and females (Data S4). This set of 932 genes was significantly enriched for sex-specific genes (Figure 5B, Table 2).

**Figure 5.**
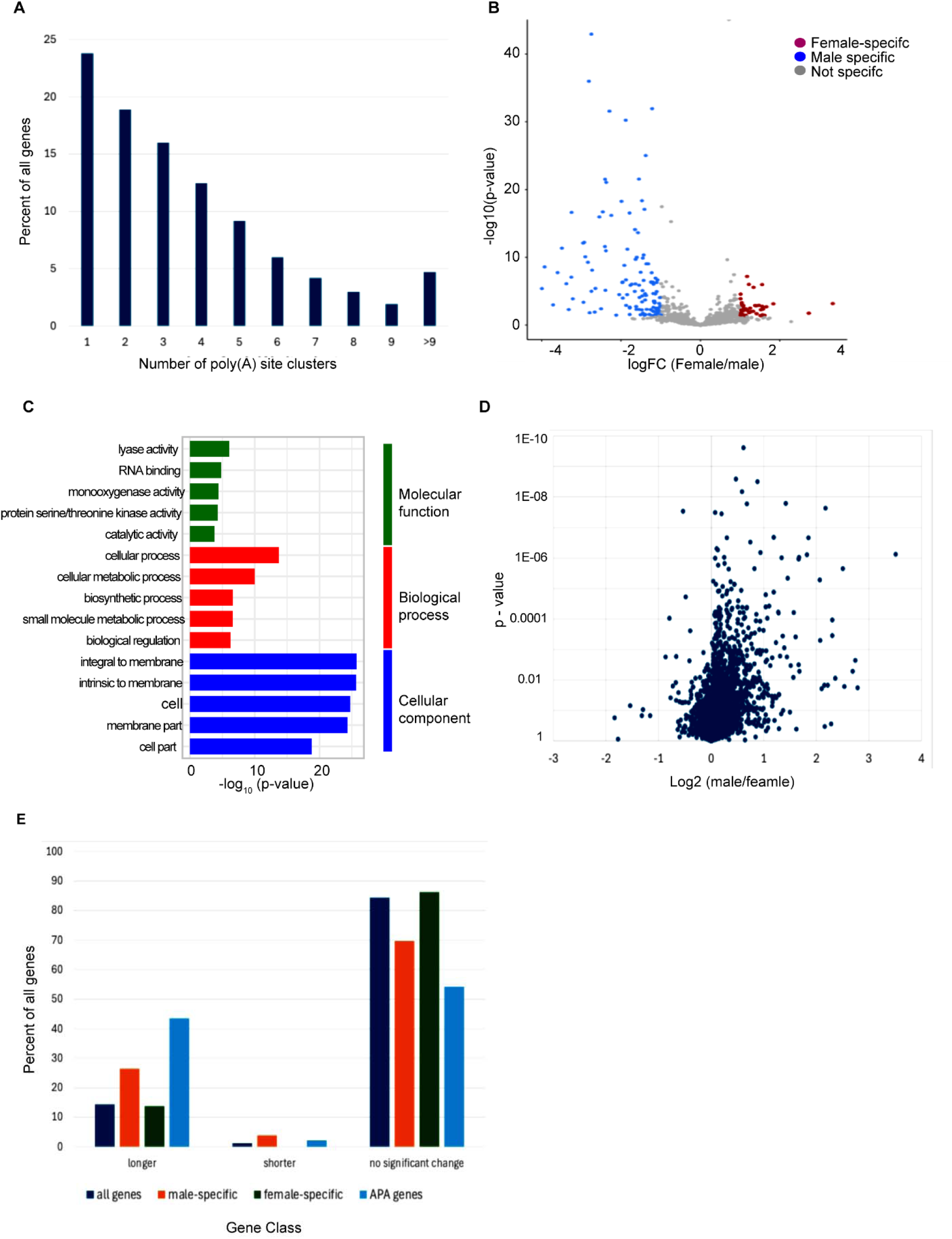
Analysis of APA genes for male and female hemp. (A) Distribution of poly (A) site clusters (PACs) per gene. A total of 69,596 PACs were identified across 18,618 genes. (B) Volcano plots showing the distribution of differentially expressed genes between male and female samples for APA genes. LOG_₂_ fold change of 2 and FDR-adjusted p-value <0.05 (C) bar plot illustrating the top 5 enriched Gene Ontology (GO) terms from each catogery (molecular function, biolofgical process and cellular compoment) for the DEGs of APA genes (D) Volcano plots showing 3□ -UTR length changes in males and females. Values on the x-axis are the log2-transformed ratios of 3□ -UTR lengths in males to females for each gene. (E) Distribution of genes based on changes in 3□ - UTR length between males and females for DEGs (□^2^ *p* value = 2.35E-13).

**Table 1.** Summary of Poly(A) site analysis.

| Particulars | Number |
| --- | --- |
| Total number of poly(A) site clusters | 69,596 |
| Total number of genes | 18,618 |
| Total number of PACs with sex-specific usage | 1,290 |
| Total number of genes with sex-specific APA | 932 |

**Table 2.** Distribution of sex-specific and non–sex-specific genes among APA-regulated and all expressed genes.

| Class | Percentage |
| --- | --- |
| **All genes** |  |
| Male specific | 671 (2.8%) |
| Female-specific | 215 (0.9%) |
| Not sex-specific | 22,457 (96.2%) |
| **APA genes** |  |
| Total | 932 |
| Male specific | 125 (13%) |
| Female-specific | 40 (4.3%) |
| Not sex-specific | 767 (82%) |
| **$X^2$ p-value = 4.06E-108** |  |

We performed a GO enrichment analysis to pinpoint the pathways enriched among the genes affected by APA (Figure 5C, Data S5). The GO terms for genes showing differential PAC usage closely resemble those from the analysis of DEGs, including RNA binding, lyase activity, catalytic activity, monooxygenase activity, and processes related to cellular, metabolic, and biosynthetic functions. These findings suggest that APA genes are predominantly associated with RNA processing, enzymatic activity, and molecular binding functions, which play a role in post-transcriptional regulation and cellular metabolism.

APA typically remodels mRNAs so that their 3□-UTRs may be shorter or longer. To assess trends in males and females, the changes in average 3□-UTR lengths for 3□ -UTRs annotated in the hemp genome annotation were measured. There was a general trend towards longer 3□ -UTRs in expressed genes in males (Figure 5D and 6E, Figure S2), a trend that was more pronounced in genes that showed higher expression in males than in females (Figure 5E). Interestingly, most of the 3□ -UTR remodeling that is seen in genes affected by APA involved 3□ -UTR lengthening (Figure 5E).

**Figure 6.**
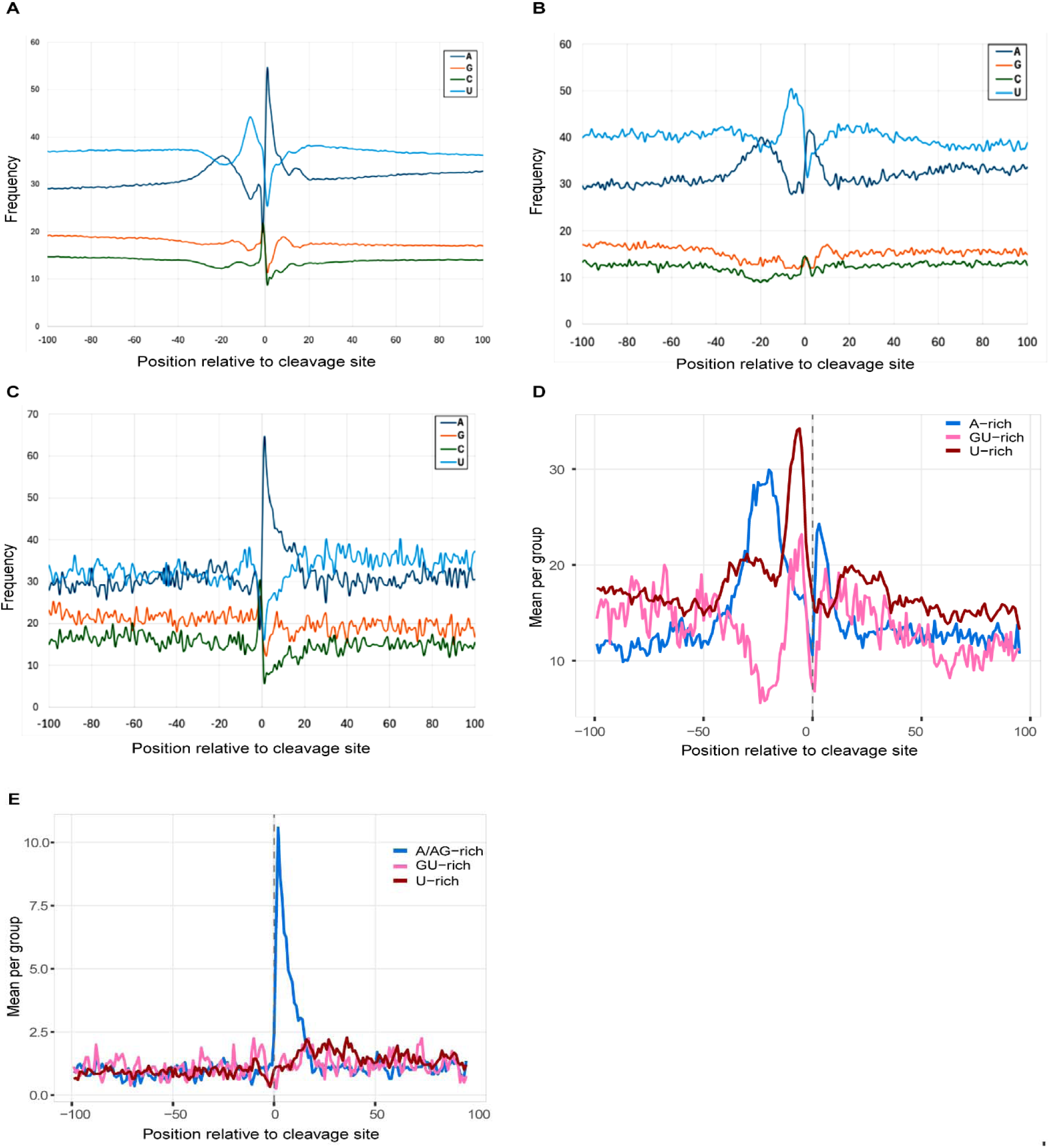
Poly(A) site usage and dynamics in male and female hemp. (A) Relative nucleotide frequency around poly(A) cleavage sites for (F) all PASs (n=460,034), (B) male-specific sites (n= 5384) and (C) female-specific sites (n=744), showing distinct nucleotide preferences near the cleavage site. (D–E) Enrichment of canonical and variant polyadenylation signal motifs upstream of (D) male and (E) female PAS.

In plants, poly(A) sites are associated with a set of three characteristic sequence signals: the Far-Upstream Element (FUE), Near-Upstream Element (NUE), and Cleavage Element (CE). These elements are defined by degenerate motifs: UGUA-related for FUE, AAUAAA-related for NUEs, and a U-rich region that flanks a YA dinucleotide for CEs. These trends can be visualized by measuring the position-by-position nucleotide frequencies of individual poly(A) site collections. The “normal” frequencies for plant genes were reflected in the trends observed for a collection of approximately 460,034 individual hemp poly(A) sites (Figure 6A). A similar distribution was observed for male-specific poly (A) sites (Figure 6B). In contrast, there were no trends in female-specific sites that reflected the canonical FUE-NUE-CE organization. We observed an increased frequency of G and C and a decreased frequency of U, contrasting with the canonical organization (Figure 6C). What is apparent in female-specific sites is a decided A-richness adjacent to the poly(A) site.

The trends in male- and female-specific sites are reflected in the distribution of abundant sequence motifs. In male-specific sites, A-rich motifs related to AAUAAA were abundant at the position corresponding to NUE (Figure 6D). In female-specific sites, GAAGAA-related motifs were abundant at the position defined by the novel A-rich peak adjacent to the poly(A) site (Figure 6E). Collectively, these results reveal a wide-ranging remodeling of hemp mRNAs by polyadenylation, with distal poly(A) sites being preferred in males. They also indicated that female-specific poly(A) sites have a novel sequence dependence that differs from male-specific sites and poly(A) sites in general.

## Discussion

### Implications of the patterns of male- and female- specific gene expression

Hemp (*Cannabis sativa*) displays a predominantly dioecious breeding pattern, with females and males having XX and XY heterogametic chromosomal arrangements, respectively (Riera-Begue et al., 2025). Important cannabinoids, including THC and CBD, are preferentially found in the glandular trichomes on inflorescence bracts in female plants. In addition, unfertilized female inflorescences produce higher levels of cannabinoids than fertilized inflorescences. Thus, across multiple hemp value chains, such as grain, and CBD production, female plants are highly desirable, and male plants are often removed to safeguard yield and quality (Toth et al., 2020). Hence, understanding the molecular regulation of sex expression and sex differentiation in female and male hemp plants becomes paramount to development of methods to distinguish female plants at early growth stages or to develop female-only hemp lines is of significant economic importance to the hemp industry.

The identification of DEGs between the two sexes at the early vegetative stage in hemp in the current study corroborates previous findings in other dioecious species, that it is possible to identify genes whose expression differs between sexes, even at the early vegetative stage. In the dioecious plant *Silene latifoli,* gene expression analysis identified two male-specific and one female-specific ESTs whose expression differed at the rosette stage long before flowering initiation (Zluvova et al., 2010). Similar findings have also been reported in *Pistacia chinensis*, where proteomic analysis identified differentially accumulated proteins between male and female plants in non-reproductive tissues at early vegetative growth Similarly, in spinach (*Spinacia oleracea*), an early floral transcriptome analysis revealed possible sex-associated DEGs, many of which map within or near the sex-determining region (Ma et al., 2022b). Even in hemp, recent transcriptome profiling of vegetative and early reproductive stages has detected sex-specific expression among these stages (Shi et al., 2025).

We identified many more male-specific differentially expressed genes (DEGs) than female-specific DEGs in this study. This finding aligns with other studies conducted on *Asparagus officinalis* (Harkess et al., 2015) and *Salix viminalis* (Darolti et al., 2018), which also reported a higher number of male-specific genes than female-specific ones. This predominance of male-specific genes is hypothesized to reflect the evolutionary development of dioecy plants, which are characterized by a more rapid divergence in males (Muyle, 2019, Zemp et al., 2016). Among the most abundant male-specific transcripts, several genes were implicated in RNA binding, monooxygenase activity, and cellular and metabolic activity. We observed the highest fold change of 94 in males compared with females for the gene CsFN_00G0021400, which encodes histone H3-K9 methyltransferase. Another gene, CsFN_00G0037350 (histone lysine-specific demethylase), also showed high expression in males, with a fold change of 42 (Data S2). These genes indicate that epigenetic regulation and chromatin modification might influence sex-specific gene expression. Similar observations of sex-specific DNA methylation-related genes have been reported for Populus balsamifera (Bräutigam et al., 2017), papaya (Zhou et al., 2020), and andromonoecious poplar (Song et al., 2012). Moreover, several thioredoxin-related genes were upregulated in males compared with females in our study, with one gene showing an 87-fold change (CsFN_00G0004070) and others exhibiting a 2-fold change (CsFN_04G0018640, CsFN_06G0033720, CsFN_10G0004900, CsFN_07G0013900, CsFN_09G0024100). In *Arabidopsis thaliana*, thioredoxin reductase, which is highly expressed in pollen grains, is known to contribute to male flower development by maintaining thioredoxin in its active reduced state. A reduction in thioredoxin reductase activity has been associated with defective pollen development and pollen lethality. Therefore, the higher expression of thioredoxin-related genes in males may indicate a role for redox regulation in male reproductive development (Reichheld et al., 2007, Marty et al., 2009, Traverso et al., 2013).

Sex conversion in plants has been documented through the use of plant growth regulators like ethephon and silver thiosulfate. These treatments influence hormone pathways, particularly ethylene, and can induce sex reversion by altering floral development programs in hemp, hops (*Humulus*), and cucurbits (Monthony et al., 2026, Akagi et al., 2025, Zhang et al., 2017a). Consistent with this, several ethylene-related genes, including ACC synthase, ACC oxidase, and ethylene response factor (ERF) genes, have been reported as differentially expressed during sex reversion in hemp (Monthony et al., 2024, Monthony et al., 2026, Garcia-de Heer et al., 2025). *ERF1* (LOC115723050) has been implicated in the ethylene signaling pathway and is highly upregulated in induced male flowers (produced by silver thiosulfate treatment of female plants) than in female flowers (Monthony et al., 2026). In line with this pattern, putative homologs of *ERF1*, such as CsFN_07G0024990, CsFN_06G0014730, and CsFN_06G0014740, also show upregulation in male flowers in our study, each exhibiting an approximately two-fold increase in expression. More broadly, members of the AP2/ERF transcription factor family are known to regulate floral organ identity and development in model species such as Arabidopsis and rice (Teng et al., 2025, Bowman et al., 1991). These findings suggest that ERF-family transcription factors may play a role in flower development. Another hormone, gibberellins (GA) have also been linked to sex expression and the promotion of male flowers (Zhang et al., 2017a, Zhang et al., 2017b, Ram and Jaiswal, 1972). In our study, we detected strong differential expression of a GA-associated gene, CsFN_00G0049550, annotated as a gibberellin-regulated protein, which showed a 19-fold higher expression in male flowers compared with female flowers. Taken together, our ethylene- and GA-related expression patterns suggest that hormone-regulated transcriptional networks responsible for floral induction are already active in the early vegetative stage.

The timing of sex-specific gene expression in plants is not well understood, and existing studies report considerable variation among species. In the dioecious annual herb *Mercurialis annua*, sex-specific gene expression begins early during plant development. Sex-specific gene expression was detected at the first-leaf stage and peaked before the onset of flowering (Cossard et al., 2019). In contrast, studies on *Silene latifolia* have indicated that sex-specific expression becomes evident later, during the rosette stage that precedes flowering (Zluvova et al., 2010). In hemp, morphological differentiation between male and female plants can be observed as early as the four-leaf stage (Shi et al., 2025). Consistent with this early phenotypic differentiation, the detection of sex-specific gene expression in four-day-old seedlings in our study was not unexpected (Figure 3 and 4A). Notably, we were able to show that sex-specific genes were consistent across genotypes, suggesting that early sex-specific expression is a conserved feature. Our data indicated that sex-specific gene expression at the seed stage was variable and lacked a consistent pattern (Figure 4B). This variability suggests that sex-specific transcriptional programs are not established during seed development but may instead be set and stabilized during germination or early plant establishment. Further temporal and tissue-specific analyses during seed development, maturation, and germination promise to provide new insights into the mechanisms by which male- and female- specific developmental programs are established.

### Alternative polyadenylation contributes to the sex-specific gene expression program in hemp

APA in plants has been linked to specific developmental processes, including seed germination (Cyrek et al., 2016, Fedak et al., 2016, Li et al., 2023), flower development (Zhang et al., 2015, Simpson et al., 2003, Hornyik et al., 2010, Liu et al., 2010, Czesnick and Lenhard, 2016), and root development (Conesa et al., 2020, Liu et al., 2014, Tellez-Robledo et al., 2019). In Spinach (*Spinacia oleracea*), genome-wide analyses have also shown that many genes exhibit both alternative polyadenylation and alternative splicing specific to male and female plants (Li et al., 2020). Our APA analysis in hemp showed that 76% of genes possess more than one poly(A) site, and APA was a distinguishing feature of genes expressed differentially in male and female hemp plants. In particular, a switch from proximal to distal poly(A) sites is disproportionately observed in genes whose expression is greater in males than in females (Figure 5D and 5E). Conversely, there was not a noticeable change in proximal vs distal site preference in female-specific genes (Figure 5E). Although a direct cause-and-effect relationship was not addressed in this study, these results suggests that an important part of the sex-specific gene expression program involves shifts between the usage of proximal and distal poly(A) sites in males and females.

Male-specific poly(A) sites have a nucleotide distribution typical of plant poly(A) sites, including sites utilized in both males and females (Figure 6A and 6B). They are also associated with AAUAAA-related motifs in the position of the NUE (Figure 6D), indicating that male-specific sites are not distinguished by alternate poly(A) signal motifs. The apparent lack of distinguishable features in male-specific sites compared with “common” sites raises questions concerning the male-specific use of such sites. Very subtle motifs not identifiable with the motif-identifying tools used here may work in concert with additional accessory factors to promote the appropriate usage in development of male-specific sites. Alternatively, these hypothetical accessory factors may inhibit the use of male-specific sites in females.

In contrast, female-specific sites had a novel nucleotide distribution and were associated with possible AG-rich motifs (Figure. 6C and 6E). As with male-specific sites, this novel motif may be a positive determinant that promotes the use of these sites in females, or it may be a negative determinant that inhibits the use of these sites in males. The nature of this novel motif raises interesting possibilities. FPA, a well-characterized RNA-binding protein in Arabidopsis, is a key component of the 3□ end processing machinery that promotes proximal poly(A) site usage (Cao et al., 2025, Parker et al., 2021). Loss of FPA function leads to distal poly(A) site usage and the production of longer transcripts (Cao et al., 2025, Hornyik et al., 2010, Lyons et al., 2013, Parker et al., 2021). FPA functions in concert with the Arabidopsis orthologs of PCF11 and CLP1 (PCFS4 and CLPS3, respectively) to recognize GA-rich sequences downstream of the cleavage or polyadenylation site to promote proximal poly(A) site usage (Cao et al., 2025). The features of female-specific sites described in this study, especially their association with AG-rich sequences that lie immediately downstream of the sites, are similar to the features of proximal sites in *Arabidopsis* whose usage is mediated by the FPA-PCFS-CLPS complex. Perusal of hemp genome sequences revealed a Cannabis sativa cv Finola gene, CsFN_08G0018410, annotated as a splicing-factor-like protein homologous to Arabidopsis FPA. This gene is upregulated in females compared to male tissues (Figure S3). Similarly, another study on Hemp has shown the SPOC_FPA-like RRM_SF superfamily gene (LOC115713521) as one of the hub genes in the weighted gene co-expression network (WGCNA) network associated with female sex development (Shi et al., 2025). FPA homologues showing this differential expression pattern support the idea that hemp FPA may promote the increased use of proximal poly(A) sites enriched for AG-rich motifs. FPA proteins contain an RNA-recognition motif (RRM) and a SPOC domain, both of which are key for controlling RNA 3□ -end formation in flowering plants. In Arabidopsis, these domains allow FPA to regulate flowering time independently of day length by directing the selection of proximal poly(A) sites. The presence and activity of an FPA-like gene in hemp that favors proximal site selection raises the possibility that FPA contributes to sex-specific RNA processing and is one of the regulatory factors shaping early sex differentiation.

To summarize, male-specific transcripts were detectable as early as 4-day-old seedlings but the earliest stage at which sex-specific expression is established is still unclear because male-specific transcripts showed variable detection in seeds (Figure 7). In parallel, our model also proposes that sex differences involve post-transcriptional regulation through APA. Male-specific genes preferentially used distal poly(A) sites and were associated with canonical NUE motifs, consistent with efficient 3□ end processing and the production of longer 3□ UTR isoforms that may add regulatory capacity. In contrast, female-specific genes showed greater use of proximal poly(A) sites and contained distinctive A-rich regions near poly(A) sites, suggesting a different mode of poly(A) site recognition and a tendency toward shorter 3□ UTR isoforms (Figure 7). Together, these observations support a framework in which early sex-linked transcriptional signals and sex-biased APA jointly shape gene expression differences between hemp males and females.

**Figure 7.**
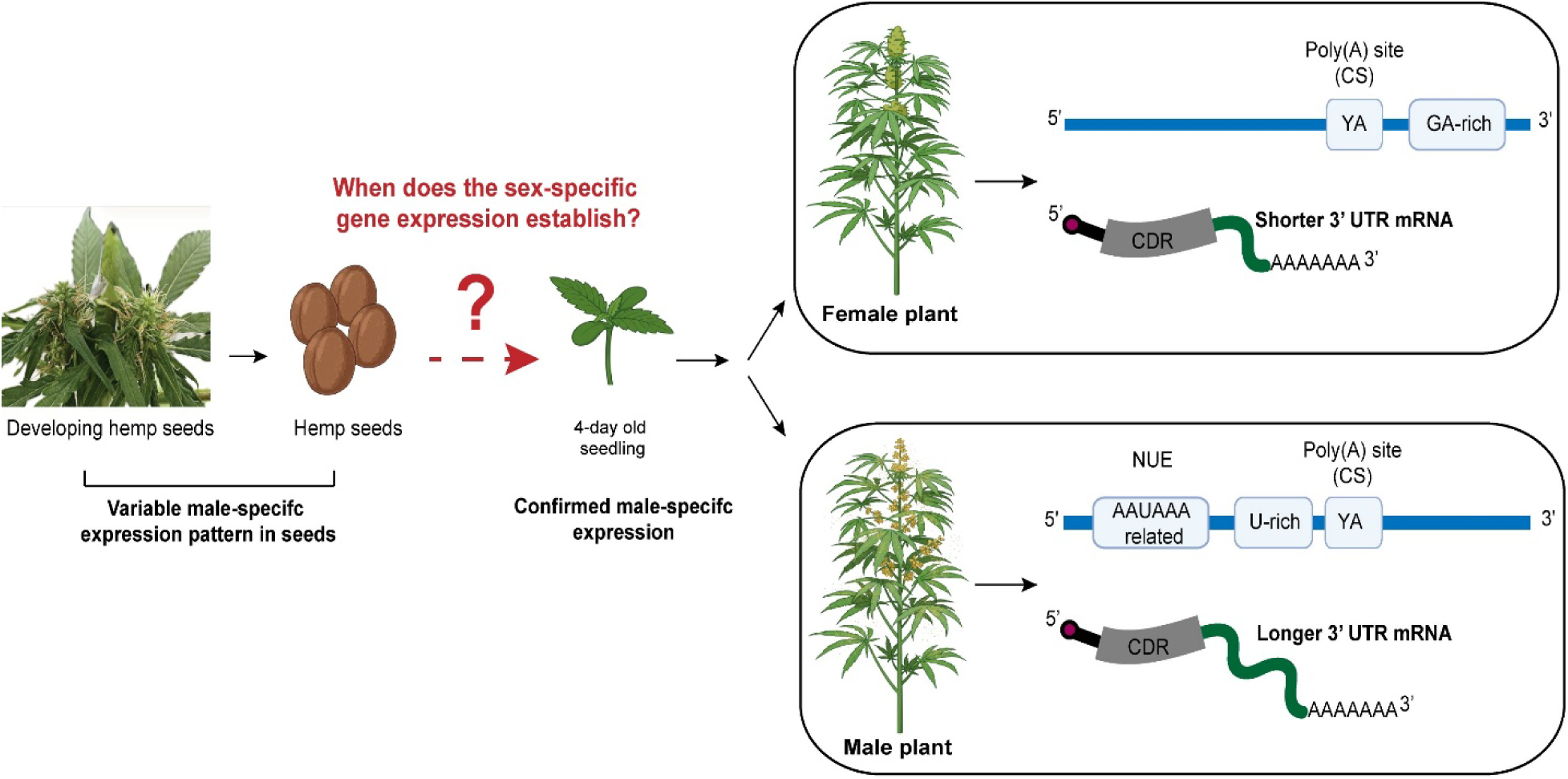
Model for sex-specific gene expression and differential alternative polyadenylation (APA) in hemp males and females. Male-specific genes can be detected very early in development, with diagnostic RT-PCR identifying male-specific transcripts in 4-day-old seedlings. The earliest stage at which sex-specific expression is established remains unclear because male-specific transcripts show variable expression in seeds. The model proposes sex-associated differences in APA and RNA processing. Male-specific genes preferentially use distal poly(A) sites and are associated with canonical near-upstream element (NUE) motifs, whereas female-specific genes show greater usage of proximal poly(A) sites and contain distinctive A-rich sequence regions near poly(A) sites. The blue horizontal line represents the proposed DNA region extending from the 5′ end to the 3′ end. The light blue boxed element labeled “YA” marks the poly(A) site / cleavage site (CS), and the other light blue boxes indicates motifs for male and female hemp plants. The magenta circle at the 5′ end denotes the 5′ terminus or cap region of mRNA transcript. The gray block labeled CDR represents the coding region. The green curved segment represents the different length of 3′ untranslated region (3′ UTR) for male and female hemp plants generated after cleavage and polyadenylation at the proximal poly(A) site. The “AAAAAAA” sequence at the 3′ end indicates the poly(A) tail.

## Conclusion

A comprehensive transcriptome of early vegetative tissues of hemp plants of both sexes was generated. Our analysis identified a set of genes that showed both differential expression and APA between the two sexes at early vegetative stages. Male-specific genes were identified as early as in 4-day seedlings way before the onset of reproductive development and were expressed across genotypes. However, in seeds, the sex-specific gene expression program was not apparent. Male-specific genes tended to use distal poly(A) sites with canonical NUE elements, whereas female-specific genes showed greater use of proximal poly(A) sites with unique A-rich regions. These results point to a developmental window between germination and early vegetative growth as the time during which sex-specific gene expression is established. They also implicate APA as a key regulatory mechanism in establishing this program. These results provide a strong foundation for developing reliable molecular tools to distinguish male plants early in the cultivation process.

## Materials and Methods

### Plant material and sample collection

The experiment was conducted in a greenhouse at the University of Kentucky. The PRO-MIX-BX growing medium (Premier Horticulture Inc.) was used as the potting mix, and Peters professional (20 N: 10 P_2_O_5_:20 K_2_O) was used for fertilization. The seeds were sown and germinated in tray pots. After 7 days of sowing, the seedlings were transferred to 5 kg pots containing the growing medium. Fertilizer water was provided twice a day for the duration of the crop.

For sample collection of RNA library preparation, 40 plants of *Cannabis sativa* cv. ‘Finola’ was grown. Each seedling was labeled, and leaf tissue samples were collected from hemp seedlings that were seven days old. Tissue samples were collected at the same time during the day to avoid any diurnal variation, and upon collection, the tissue samples were flash frozen in liquid nitrogen and stored at −80°C for further study. Hemp plants were grown until the onset of reproductive structures to phenotypically determine the sex of individual plants.

For sample collection of RT-PCR experiment, plants of *Cannabis sativa* cv. ‘X-59’ was grown. Throughout the experiment, the photoperiod was consistently set to 18 hours of daylight and 6 hours of night, as the flowers are not sensitive to changes in photoperiod. Each plant was labeled, and leaf tissue samples were collected from hemp plants at the ages of 4, 7, 14, 24, and 28 days after sowing. The samples were gathered at the same time each day, and immediately after collection, they were flash-frozen in liquid nitrogen and stored at −80°C for further study.

### RNA extraction and RNA library preparation

Total RNA was extracted using Ambion TRIzol (Life Technologies) and precipitated with ethanol. 50-100 mg of collected leaves or individual seeds were crushed with liquid nitrogen, and TRIzol was used for nucleic acid extraction. RNA was purified using RNA purification columns (Enzymax LLC) and Qiagen reagents, following the manufacturer’s recommendations. The final RNA concentrations ranged from 50-2,000 ng/μL.

We used a 3□ -end-directed transcriptome approach (also referred as PATseq or Poly(A) Tag Seq), where short cDNA tags that correspond only to 3□ -ends of transcripts were prepared, sequenced and quantified (de Lorenzo et al., 2017, Chakrabarti et al., 2018). The RNA was diluted to 2-10ug for PATseq library preparation and were prepared as described previously (Zhou and Li, 2023, Ma et al., 2014). Total RNA was fragmented for 2 min at 95°C with 10X RNA Fragmentation buffer (New England Biolabs-NEB) and eluted as described by the manufacturer. This reaction can be empirically tuned to generate a small number of random cleavage events in a population of RNAs. The fragmented RNA was mixed with magnetic beads with immobilized oligo-dT (NEB) for poly(A) enrichment. This method selects affinity-purifying poly(A)-containing fragmented RNAs using immobilized oligo-dT. Magnetic beads were collected and washed with washing buffer B (10mM Tris-HCl PH 7.5, 0.15M LiCl, 1mM EDTA). Further 10 mM Tris-HCl was added and incubated at 80□C and for 2 min, followed by incubation with binding buffer (NEB) for the second round of poly(A) enrichment. Subsequently, the RNA was eluted using nuclease-free water. PolyA-enriched RNA samples were used for cDNA synthesis using Smartscribe (Takara). The reaction mixture (25 µL) was prepared with 1X first-strand buffer, 10 mM dNTPs, 1 mM DTT, 100 pmol RT primer (Data S1), and 1 µL of enzyme. After 60 minutes at 42°C, 100 pmol of the strand-switching primer (SMART7.5) and an additional 1 µL of enzyme were introduced, and the reactions were incubated for another 60 minutes at 42°C. After a final incubation at 70°C for 5 min to stop the reaction, SPRI beads (HighPrep PCR, Magbio Genomics, Inc.) were used to purify the solution. The SPRI beads incubated with cDNA was incubated for 8 minutes at room temperature were then washed with 80% ethanol, air-dried, and cDNA was eluted with 25 µL water. 1µl of the eluted cDNA was used for PCR amplification using Phire Hot Start II DNA Polymerase (Thermo Fisher Scientific) with primers PE-PCR1 and PE-PCR2 (Data S1). The cycle temperatures and durations were 95°C for 15 seconds, 60 °C for 15 seconds, and 72°C for 60 seconds. The reactions were performed for 15 cycles. PCR products were separated on 1% agarose gels, and products of approximately 500 bp were excised and purified using a Qiagen gel purification kit. The gel-purified fragments were re-amplified using the same PCR conditions and cleaned using SPRI beads. This final library was quantified using a Qubit and submitted for sequencing on an Illumina instrument at the University of Kentucky HealthCare Genomics Core Laboratory, USA.

### Transcriptome data analysis

PATseq libraries were sequenced on the Illumina platform. Raw data were imported and processed using CLC Genomics Workbench (version 23). The raw sequencing data were mapped to a customized set of sequences consisting of organellar and ribosomal RNA sequences and unmapped reads retrieved for further analysis; the mapped reads were discarded. The unmapped reads were demultiplexed to assign reads to individual samples based on unique barcodes. Additionally, the reads were trimmed to remove poly(A) tails and Illumina adapter sequences and further trimmed at their 3□ ends to a length of 40 nts. Processed reads were aligned to the *Cannabis sativa* cv ‘Finola’ genome sequence (SuperCann database: https://gdb.supercann.net/index.php/download) using the “Map Reads to Sequence” tool available in the CLC Genomics Workbench. For mapping PATseq reads to the Finola genome, the following parameters were used: mismatch cost-2, insertion cost-3, deletion cost-3, length fraction-0.9, and similarity fraction-0.9. The mapping results were exported as bam files for further processing. Summaries concerning the reads and mappings are provided in Table S1.

Bam files were subsequently converted to the bed format using the bamtobed tool in the Bedtools suite. Mapped reads were converted to single-nucleotide reads and re-mapped using the annotateBed tool to a modified hemp genome in which all annotated genes had been extended at their 3□ ends by 500 bp. The results were then ported in CLC Genomics Workbench for further analysis using the Microarray Analysis set of tools. Briefly, genes with no mapped reads or genes with expression across all samples (male and female) were 0 were removed, the raw expression values (mapped reads) were transformed by adding 1 to every value, and the transformed expression values normalized to yield final expression values in tags per million. Baggerley’s test, a proportion-based statistical test, was conducted to identify genes displaying significant differences in expression between the two sexes. Genes showing more than 2-fold change in expression and an FDR-corrected p-value of <0.05 between the two sexes were defined as differentially expressed genes (DEGs).

For Principal Component Analysis (PCA), the entire set of expressed genes (23,343 genes) was used. A gene was considered expressed if it showed a non-zero expression value in at least one sample. For hierarchical clustering, only DEGs were used, and Euclidean distance and average linkage were used as distance metrics and linkage methods for visualizing sample-relatedness and DEGs co-variation. Enrichment of gene ontology terms was performed using singular enrichment analysis (SEA) in AgriGO v2.0. We used Fisher’s exact test with FDR correction (cutoff <0.05), “complete GO” ontology, a minimum term size of >5 mapped genes. The data were used to generate figures showing the top 5 GO terms within each category using the ggplot2 package in RStudio 4.1.

### Alternative poly (A) site and 3□ -UTR analysis

To analyze poly(A) site usage between male and female samples, trimmed (single nucleotide) PATseq reads were mapped using annotateBed tool in BedTools to a modified Finola genome dataset in which annotated 3□ -UTRs were extended by 500bp. For each gene, the average 3□ -UTR length was calculated as the weighted sum of the relative lengths of individual poly(A) sites using the following formula:

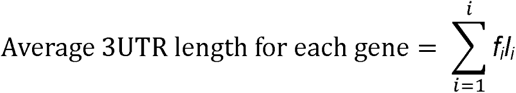

where *f*_₁_ represents the fractional usage of each poly(A) site (with values between 0 and 1) and *l_1_* is the relative length with respect to the extended annotated 3□ -UTR, with reads extending to the end of the 3□ -UTR having a value of 1. The ratio of average 3□ -UTR lengths between male and female samples was computed, and log_₂_-transformed values were plotted to assess the genome-wide differences in 3□ -UTR length distribution between the sexes.

For the nucleotide composition around polyadenylation sites, individual sites were classified as male-specific, female-specific, or common. Changes in poly(A) site usage were determined using the DEXSeq package (Anders et al., 2012) as described previously (Chakrabarti et al., 2020). Sex-specific sites were defined as those showing a 10-fold or greater relative usage in males or females, with an FDR p-value cutoff of 0.05. A ±100 nt genomic window centered on the cleavage site (position 0) was extracted for each poly(A) site (PAS). The nucleotide frequencies (A, C, G, and U) were calculated at each position across all PASs, as well as separately for male- and female-specific sites, to visualize the sequence preferences surrounding the cleavage site. For motif discovery and positional enrichment, the region upstream of each PAS was scanned using SignalSleuth (Loke et al., 2005) to identify canonical and variant polyadenylation signal motifs.

### Confirmation of DEGs using RT-PCR

For reverse transcription, 1-2 µg of total RNA isolated from leaves and seeds were isolated using Ambion TRIzol (Life Technologies). Reverse transcription reactions were initiated with RT primers (Data S1), utilizing the Smartscribe enzyme (Takara) as the reverse transcriptase. Only the synthesis of the first-strand cDNA was conducted, following the same procedure as previously mentioned. The first-strand cDNA was then purified using SPRI beads, employing the same method described earlier. 1µl of the eluted cDNA was used for PCR amplification using Phire Hot Start II DNA Polymerase (Thermo Fisher Scientific) with selected RT-PCR primers (Data S1). The reaction mixture (25 µL) was prepared using 1X buffer, 10 mM dNTPs, and 1 µL of enzyme. The cycle temperatures and durations were 95°C for 15 seconds, 60 °C for 15 seconds, and 72°C for 60 seconds. The reactions were performed for 35 cycles.

## Acknowledgements

We thank Carol Von Lanken for lab and greenhouse support, including plant growth and maintenance. Artificial intelligence-based language correction software was used in the text to ensure accurate spelling and grammar, without influencing the content. Images were generated using BioRender (https://BioRender.com) and Adobe Illustrator software.

## Short legends for Supporting Information

| File name | Description |
| --- | --- |
| Supplemental figures | **Figure S1:** Browser tracks representing expression of selected DEGs in two sexes |
|  | **Figure S2:** Genome browser view of selected genes showing sex-specific poly(A) site usage in males and females |
|  | **Figure S3:** Normalized gene expression values of CsFN_08G0018410 gene for males and females. |
| Supplementary table | **Table S1:** Summary of read filtering, demultiplexing, and mapping for pooled PATSeq libraries |
| Supplemental File 1 | **Data S1:** RT primers information |
| Supplemental File 2 DEG | **Data S2:** Summaries of the DEG analysis |
| Supplemental File 3 | **Data S3:** GO term analysis for all differentially expressed genes |
| Supplemental File 4<br>DEXSeq | **Data S4:** Summaries of the DEXSeq analysis |
| Supplemental File 5 | **Data S5:** GO term analysis showing the differential PAC usage |

