## Supplemental Figures S1-S3 for "Alternative polyadenylation and the sex-specific gene expression program in hemp"

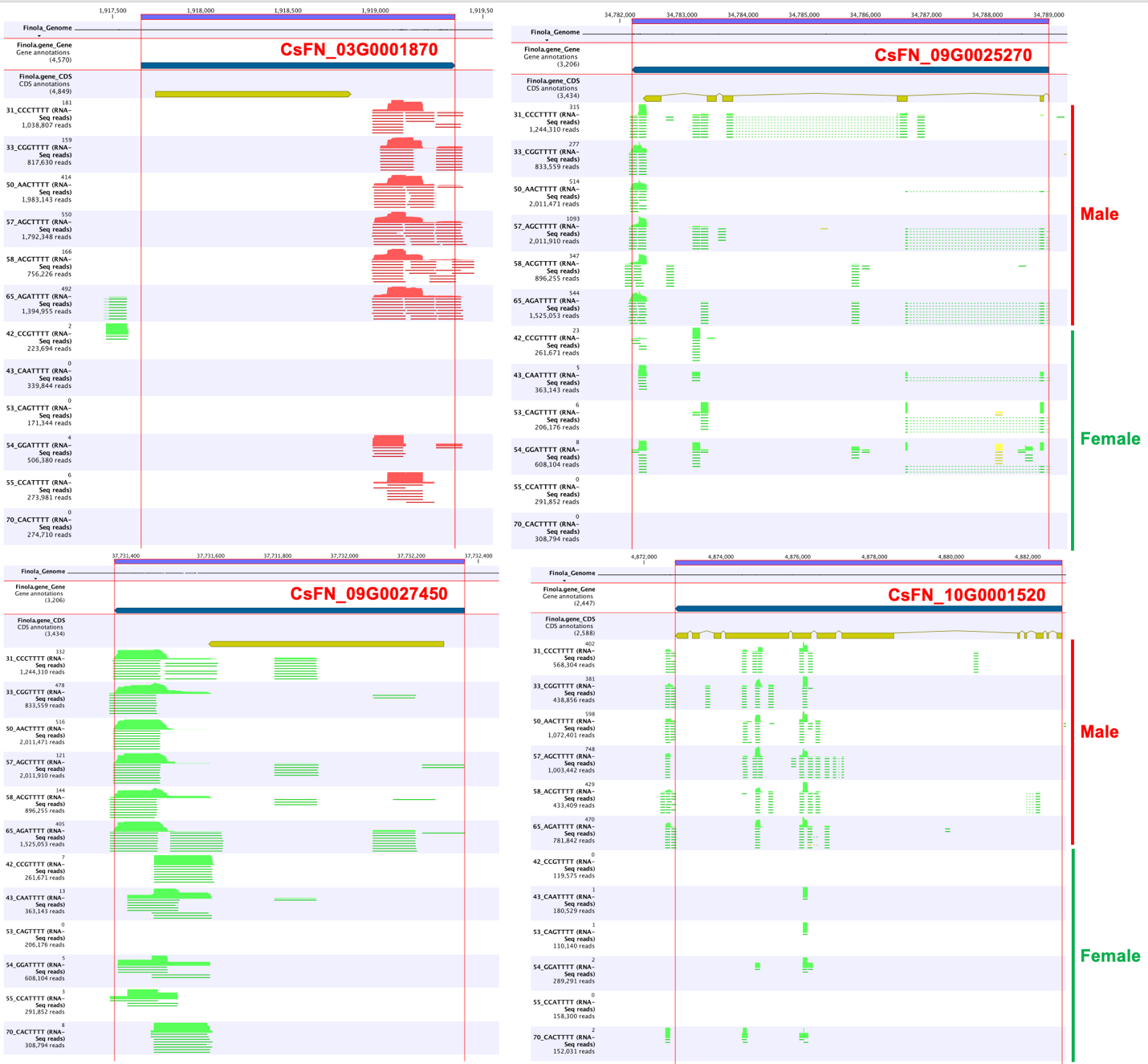

**Figure S1: Browser tracks representing expression of selected DEGs in two sexes.** Browser tracks represent mapping of PATseq reads to the respective genes in the hemp genome. Region demarcated by red vertical lines represent respective DEG. Male and female samples are marked with red and green lines, respectively.

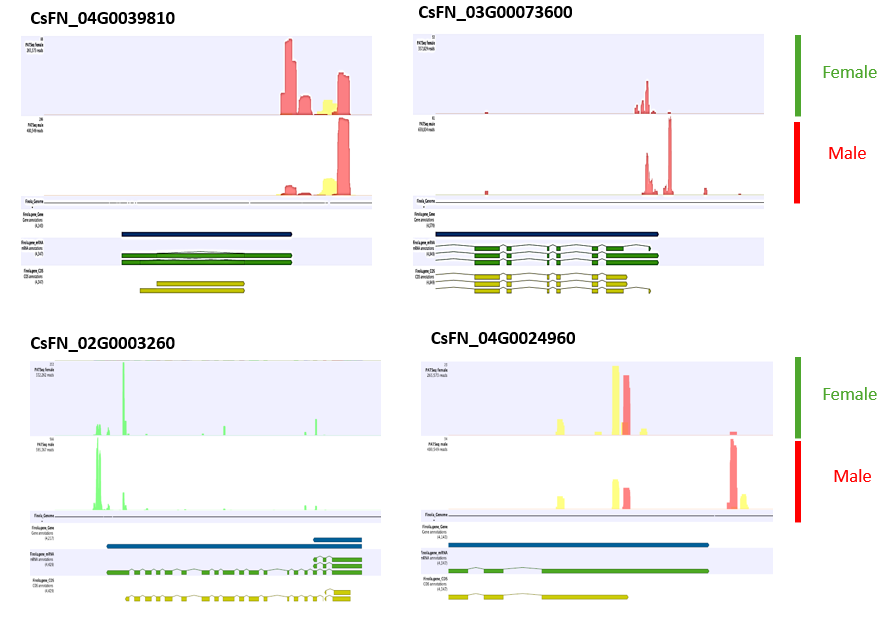

**Figure S2: Genome browser view of selected genes showing sex-specific poly(A) site usage in males and females**. Browser tracks represent mapping of PATseq reads to the respective genes in the hemp genome. Region demarcated by colored lines poly(A) site usage. Male and female samples are marked with red and green lines, respectively.

CsFN_08G0018410

| Hemp plants | Normalized values |
| --- | --- |
| Male | 16.4465 |
| Female | 13.4396 |

**Figure S3: Normalized gene expression values of CsFN_08G0018410 gene for males and females.**
