## Supplementary Table S1 for "Alternative polyadenylation and the sex-specific gene expression program in hemp"

**Table S1:** Summary of read filtering, demultiplexing, and mapping for pooled PATSeq libraries. Reads removed and remaining are shown for each processing step, with percentages calculated relative to the number of reads entering that step. Final mapping percentage represents the proportion of nuclear ‑RNA and mRNA reads successfully aligned to the reference genome.

| **Processing step** | **Reads input** | **Reads removed** | **Reads remaining** | **Percentage removed (of step input)** |
| --- | --- | --- | --- | --- |
| Total raw reads | 2696890404 | - | 2696890404 | - |
| rRNA filtering | 2696890404 | 689410505 | 2007479899 | 25.5% |
| Organelle and PhiX filtering | 2007479899 | 1410084054 | 597395845 | 70.2% |
| Demultiplexing | 597395845 | 279676490 | 317719355 | 46.8% |
| mRNA reads selection | 317719355 | 111890900 | 205828455 | 35.2% |
| **Total mapped reads (nuclear- rRNA + mRNA)** | **60.7%** | | | |
